# Humanized Bone Marrow-Liver-Thymus Mice for Studying HIV-1 Persistence in Liver and Lung CD4+ T and Myeloid Cell Subsets during Antiretroviral Therapy

**DOI:** 10.1101/2025.01.31.635552

**Authors:** Amélie Cattin, Tram NQ Pham, Jonathan Dias, Natalia Fonseca Do Rosario, Laurence Raymond Marchand, Olga Volodina, Jean-Victor Guimond, Natalie Patey, Yuanyi Li, Kathie Béland, Mohammad-Ali Jenabian, Jérôme Estaquier, Elie Haddad, Éric A. Cohen, Petronela Ancuta

## Abstract

**Background:** While the role of CD4^+^ T-cells in HIV-1 reservoir persistence during antiretroviral therapy (ART) is well-established, studies on tissue-resident macrophages (MΦ) in people with HIV-1 (PWH) are restricted by difficulties in accessing deep tissue samples. Investigations in myeloid-only humanized mouse models demonstrated the contribution of MΦ to viral rebound upon ART interruption. Two distinct MΦ subsets exist in mice and humans: one of embryonic origin and self-renewal capacity generating long-lived tissue-resident MΦ (LL-TRM), and another one of short-lived MΦ (SL-MΦ), constantly replenished by bone marrow monocytes. The relative contribution of LL-TRM *versus* SL-MΦ to tissue HIV-1 reservoir persistence during ART remains understudied. Here, we used a humanized BM-liver-thymus (hu-BLT) mouse model to quantify integrative HIV-1 infection in liver/lung MΦ *versus* CD4^+^ T-cells before/after ART and document their expression of LL-TRM/SL-MΦ markers.

**Results:** Lung/liver immune cells were extracted from ART-naive (ART-) and ART-treated (ART+) HIV-infected (HIV+) hu-BLT mice, as well as HIV-uninfected mice (HIV-). MΦ were identified as cells expressing the myeloid markers CD33/HLA-DR and/or CD68 and flow-cytometry sorted based on their differential expression of CD14 and/or CCR2. Matched CD3^+^CD4^+^ T-cells were sorted in parallel and used as controls. HIV-DNA integration was measured by nested real-time PCR. In contrast to CD4^+^ T-cells that carried the highest levels of proviral HIV-DNA before and after ART, integrative infection in liver/lung MΦ was detected before ART, but was drastically reduced in HIV+ART+ hu-BLT mice, regardless of CD14 or CCR2 expression on MΦ. Markers of LL-TRM (CD163/CX3CR1/Ki67/c-Kit) were expressed on a small fraction of liver but not lung MΦ, indicative of a deficient LL-TRM development in this hu-BLT model.

**Conclusions:** Together our results demonstrate that lung/liver MΦ in hu-BLT mice support integrative HIV-1 infection *in vivo*, but their contribution to viral reservoir persistence during ART is minor when compared to CD4^+^ T-cells. This is consistent with the deficient development of LL-TRM we observed in hu-BLT mice. However, HIV-1 permissive MΦ present in this model likely contribute to viral rebound upon ART interruption. Therefore, HIV-1 cure interventions that are tested in such preclinical models should consider targeting HIV-1 replication in both MΦ and CD4^+^ T-cells.

## BACKGROUND

Antiretroviral therapy (ART) efficiently controls human immunodeficiency virus type 1 (HIV-1) replication to undetectable levels in the plasma of people with HIV-1 (PWH) (1). However, HIV-1 is not eradicated by ART, as demonstrated by the persistence of viral reservoirs in the blood and multiple deep tissues, with viral rebound rapidly occurring upon treatment interruption (2–6). While the role of CD4^+^ T-cells in HIV reservoir persistence during ART is well-established, the contribution of myeloid cells is still controversial (7–12). This is explained by the fact that MΦ mainly reside in the peripheral tissues, while CD4^+^ T-cells recirculate in the blood, thus facilitating their non-invasive access. Nevertheless, pioneering *in situ* investigations performed in the pre-ART era demonstrated that HIV-1 efficiently targets MΦ for productive infection *in vivo* (13–15). One requirement for the persistence of viral reservoir cells is their capacity to survive long-term *in vivo*. Most recent advances in the field of myeloid cell ontogeny revealed the existence of two distinct pools of tissue MΦ: one pool of long-lived tissue resident-MΦ (LL-TRM) with self-renewal capacity, derived from embryonic stem cells of the yolk sac and fetal liver; and another pool of short-lived MΦ (SL-MΦ), derived from bone marrow (BM) monocytes under constitutive and inflammatory conditions (16–20). Among key functional differences between these myeloid cells, it is noteworthy to state that LL-TRM and SL-MΦ distinctly contribute to tissues homeostasis and inflammatory reactions, respectively (21–23). Although the presence of LL-TRM in the brain, liver, lungs, and dermis is historically well-established, recent studies demonstrated the existence of LL-TRM in multiple other tissues, including blood vessels and the heart (24). After birth, BM monocytes gradually replenish the pool of LL-TRM in specific tissues under steady-state and inflammatory conditions, and this in a CCR2-dependent manner (25, 26). Interestingly, mice studies demonstrated that in the heart, monocyte-derived SL-MΦ become predominant over LL-TRM during inflammation and aging (23). At the opposite, gut-associated lymphoid tissues (GALT) are mainly seeded by SL-MΦ, a pool constantly replenished by BM-derived monocyte (16, 18–20, 27).

Most of the current knowledge on monocyte/MΦ ontogeny emerged from mouse studies, with LL-TRM being less defined in terms of phenotype and function in humans. Different markers have been identified to discriminate between MΦ of embryonic origin *versus* BM monocyte-derived MΦ in human tissues. Among these markers, CCR2 is the receptor for CCL2 (or monocyte chemoattractant protein-1, MCP-1) and is involved in monocyte emigration from the BM and their further recruitment into peripheral tissues (25, 28). Accordingly, CCR2 expression was proposed as a marker to distinguish between embryonically derived (CCR2^−^) and monocyte-derived MΦ (CCR2^+^) in the heart (24). Other markers can be used to discriminate between MΦ subsets with embryonic *versus* BM ontogeny, with the phenotype of LL-TRM appearing to be tissue-specific (29). For example, in the lungs, alveolar MΦ exhibit an HLA-DR^+^CD64^+^CD206^+^CD169^+^ phenotype, with low CD14 expression (30–32). In the liver, Kupffer cells have been described to exhibit a CD14^+^CD68^+^CX3CR1^+^CD163^+^Tim4^+^ phenotype; they also express MARCO, HMOX1, CD206, and VCAM1 (27, 33). In the spleen, red pulp MΦ are defined by a CD68^+^CD163^+^ phenotype, with a low expression of CD14 (34). Red pulp MΦ also express the Fc gamma receptors IIa (FcγRII/CD32a) et IIIa (FcγRIII/CD16a), but unlike monocyte-derived MΦ, the expression of FcγRI/CD64 on the red pulp MΦ is not constitutive, but inducible during inflammation (34). Thus, tissue-resident MΦ with low/undetectable expression of CCR2 and/or CD14 may represent embryonically-derived LL-TRM, while monocyte-derived SL-MΦ likely exhibit a CCR2^+^ and/or CD14^+^ phenotype.

In humans, studies by our group and others revealed that in contrast to CD4^+^ T-cells, myeloid cells infiltrating the small intestine and colon, rarely harbor HIV-DNA during ART (35–37). The latter finding may be explained by the short life and rapid turnover of myeloid cells *in vivo*, consistent with the BM-derived monocyte ontogeny of the colon-infiltrating MΦ (38). The absence of replication-competent HIV-1 was also reported in liver MΦ, a fraction of them including Kupffer cells of embryonic origin (39). Nevertheless, HIV persistence was observed in hepatocytes and liver CD4^+^ T-cells of ART-treated PWH co-infected with hepatitis B virus (HBV) (40). Nevertheless, urethral MΦ were identified as cells carrying HIV-1 reservoirs in ART-treated PWH (41). Similarly, studies by our group observed the presence of HIV-DNA in alveolar MΦ of a fraction of ART-treated PWH (42). Most recently, studies provided new evidence supporting the contribution of brain microglia (a subset of LL-TRM) to the persistence of viral reservoirs in PWH upon long-term ART (43), thus raising new matters of concerns regarding the possibility to cure HIV-1 in this peculiar anatomic site in special (44, 45) and in the whole body in general (46). Other studies performed in non-human primates (NHP) models of simian immunodeficiency virus (SIV) infection demonstrated that specific subsets of infected MΦ can persist under ART (47, 48), which is consistent with MΦ ability to support productive viral infection (10, 12, 48, 49). Thus, evidence exist that MΦ from specific tissues server as long-lived viral reservoirs. Considering the life span of MΦ derived from embryonic precursors, it is highly likely that long-lived tissue-resident MΦ (LL-TRM) are more prone to contribute to HIV-1 reservoir persistence during ART and promote viral rebound upon ART interruption, compared to short-lived monocyte-derived MΦ (SL-MΦ). Given difficulties in accessing deep tissue MΦ from ART-treated PWH for mechanistic studies and drug screening, an alternative to overcome such limits is provided by humanized mouse models that can sustain a productive HIV-1 infection and allow viral reservoir persistence during ART (50–52). Of note, studies in a model of myeloid-only mice (MoM) provided key evidence that MΦ sustain HIV-1 replication in the absence of ART and demonstrated that MΦ isolated from the liver and lungs harbor HIV-DNA in ART-treated animals, suggesting their role in HIV reservoir persistence during ART (53). Nevertheless, limited knowledge exist on the development of LL-TRM and SL-MΦ in humanized mice models in the context of HIV-1 infection and ART treatment.

In this study, we employed a humanized BM-liver-thymus (hu-BLT) mouse model (54, 55), previously used by our group (56, 57), to explore HIV-1infection and viral reservoir persistence in myeloid cells identified based on their expression of classical MΦ markers (*i.e.,* HLA-DR, CD68, CD33), as well as CD14 (30–32) and CCR2 (24), markers proposed to distinguish between SL-MΦ and LL-TRM. MΦ subsets sorted from the liver and lungs of HIV-infected hu-BLT mice before and after ART were used for extensive phenotypic characterization and quantification of proviral HIV-DNA. Matched CD4^+^ T-cells served as positive controls for integrative HIV-1 infection. Our results demonstrate that CCR2^+^/CCR2^−^ MΦ in BLT mice are permissive to HIV-1 infection in the absence of ART, but negligibly carry integrated HIV-DNA in ART-treated mice. This is in contrast to CD4^+^ T-cells, which harbored relatively high levels of integrated HIV-DNA in the liver and lungs of aviremic HIV-infected ART-treated hu-BLT mice. Finally, we performed an extensive phenotypic characterization using reported markers of LL-TRM (*i.e.,* TIM4, CD163, CX3CR1, CD169, c-kit, Ki67) and found that neither CCR2^−^ nor CCR2^+^ myeloid cells from BLT mice liver and lungs exhibit a preferential CD163^+^CX3CR1^+^ LL-TRM-like phenotype, suggesting that CCR2 expression alone cannot distinguish SL-MΦ from LL-TRM in this model. It is noteworthy that a small fraction of CCR2^+^ MΦ (<8%) from the liver, but not the lungs, expressed the surrogate proliferation marker Ki67, indicative of self-renewal capacity. These results demonstrate that MΦ from the lungs and liver of HIV-infected hu-BLT mice are permissive to integrative HIV-1 infection in the absence of ART, but negligibly contribute to viral reservoir persistence during ART, consistent with their SL-MΦ phenotype. These results support the validity of using hu-BMT mouse models for mechanistic studies on integrative HIV-1 infection in both MΦ and CD4^+^ T-cells.

## METHODS

### Generation of humanized BLT mice

BLT mice (54, 55) were generated, as we previously described (56, 57). Briefly, NOD/LtSz-scid IL-2Rγc(null) (NSG) mice (Jackson Laboratory) between 6 and 10 weeks of age were conditioned with sub lethal (2.5 Gy) total-body x-ray irradiation and human fetal thymus fragments of about 1mm^3^ (1 per mouse) were implanted underneath the recipient’s kidney capsule. Within 24 hours, 3-5×10^5^ human autologous fetal liver CD34^+^ cells in 100 µl of dPBS^−/-^ were injected intravenously. Batches of BLT mice made with fetal tissues from five distinct donors (A-E, as indicated in Figure legends) were included in this study. The efficiency of humanization in BLT mice was monitored by flow cytometry analysis of human immune cell subsets from peripheral blood drawn from mice at different weeks post-humanization. Briefly, the presence/frequency of cells positive upon staining with human (h) CD45, hCD19, hCD14 and hCD3 Abs, and negative for mouse (m) CD45 Abs, were monitored in the blood of hu-BLT mice from week 4 post-humanization to the time of infection. All animal experiments were performed upon approval by the IRCM Animal Care Committee (IRCM 2018-11) and by the CHU Sainte-Justine Animal Care and Use Committee (CIBPAR#2021-2961), respecting the Good Laboratory Practices for Animal Research and the relevant regulatory standards in accordance with institutional and national guidelines. Hu-BLT mice were generated at the CHU Sainte Justine, while HIV-1 infection, ART treatment, and sacrifice were performed in the Biosafety Laboratory Level 3 (BSL-3) at the IRCM. Subsequent immunological and virological experiments on tissue-infiltrating immune cells of hu-BLT mice were performed in the BSL-3 at the CRCHUM.

### HIV infection and ART treatment of BLT mice

BLT mice were infected with the CCR5-tropic NL4.3-ADA-GFP HIV-1 virus (200,000 TCID_50_) *via* intraperitoneal injection. HIV-1 viral load was determined in the plasma of hu-BLT mice weekly using the quantitative COBAS AmpliPrep/COBAS TaqMan HIV-1 test, version 2.0, Roche (detection limit, <20 copies/ml) (57). We previously showed that this HIV-1 strain was infectious *in vivo* (57, 58), and was appropriate to study the establishment and maintenance of HIV-1 reservoirs in hu-BLT mice (57, 59).The ART treatment, consisting of emtricitabine (100 mg/kg), tenofovir (50 mg/kg), and raltegravir (68 mg/kg), was administered daily *via* subcutaneous injections in HIV-infected hu-BLT mice starting at 6-7 weeks post-infection (wpi). The treatment was administered for a period of 4-6 weeks before the sacrifice (identified as HIV^+^ART^+^). In certain experiments, HIV-infected hu-BLT mice were not treated with ART for the entire duration of the experiment (identified as HIV^+^ART^−^).

### Tissue processing

Liver and lung cells from HIV^−^, HIV^+^ART^−^ or HIV^+^ART^+^ hu-BLT mice were extracted using mechanical and enzymatic disruption as previously described (57, 59). Briefly, tissues were cut into small pieces and digested with type II collagenase (0.025 IU/mL, Worthington Biochemical) and benzonase (10 IU/mL, EMD Millipore) for 1h at 37 °C. Tissues were then mechanically disrupted and filtered on a 70-μm cell strainer. For liver tissues, hepatocytes were removed by two subsequent slow centrifugations (50g, 5 min). Leukocytes were purified by Ficoll density centrifugation. Cells were directly stained with Abs for flow cytometry analysis or preserved in liquid nitrogen for subsequent FACS sorting.

### Flow cytometry analysis

Blood cells from hu-BLT mice obtained at different weeks post-humanization were analyzed for human cell reconstitution. Upon sacrifice, the phenotype of cells extracted from different tissues (liver, lungs) was assessed by flow cytometry. Briefly, cells were stained on the surface with a cocktail of fluorescence-conjugated Abs as listed in Supplemental Table 1. The viability dye LIVE/DEAD® Fixable Aqua Dead Cell Stain Kit (Invitrogen) was used to exclude dead cells from the analysis. In applicable experiments, surface-stained cells were then fixed and permeabilized using the BD Cytofix/Cytoperm kit (BD Biosciences) and intracellularly stained with Abs against intracellularly expressed markers as listed in Supplemental Table 1. Cells were analyzed using a Fortessa cytometer (Diva, BD Biosciences) and analyzed using the FlowJo (version 10.6.1, Tree Star Inc.).

### Flow cytometry sorting

CD4^+^ T-cells and CCR2^+^ *versus* CCR2-myeloid cells were sorted by flow cytometry (BD AriaIII). Briefly, cells extracted from the liver and lungs were stained with a cocktail of human fluorescence conjugated antibodies recognizing surface (CD3 PB, CD4 AF700, HLA-DR BV786, CD19 FITC, CD56 FITC, CD33 PE, CCR2 PE-DAZZLE 594, CD14 APC) and intracellular (CD68 PE-Cy7) markers, as well as mouse CD45 PerCP-Cy5.5 Abs, using the sorting buffer (PBS 1X with 5% FBS and 25 mM Hepes buffer). A viability dye was used to exclude dead cells from the analysis. The cell purity upon sort was >99% (Supplemental Figure 2).

### HIV-DNA quantification

Integrated HIV-DNA quantification was performed on cell lysates by nested real-time PCR (LightCycler 480, Roche) using specific HIV-LTR and Alu primers, as we previously described (36, 60). The HIV copy numbers were normalized to cell numbers upon simultaneous amplification of the CD3 gene with specific primers (two CD3 copies/cell). ACH2 cells containing one copy of proviral HIV-DNA were used to generate a standard curve (30,000; 3,000; 300, 30, and 3 cells/reaction). The limit of detection of the assay was 3 copies/test for both HIV and CD3. Each PCR was performed in triplicates using 30,000-100,000 cells/reaction.

### Statistical analysis

Statistical analysis was performed using the GraphPad Prism 8 software. The statistical tests applied are specified in Figure legends.

## RESULTS

### Generation of hu-BLT mice

Here, we used a hu-BLT mouse model, as previously reported by our group (56, 57). The objective was to investigate integrative HIV-1 infection in different subsets of MΦ *versus* CD4^+^ T-cells extracted from the liver and lungs of HIV-infected ART-naive (HIV^+^ART^−^) and ART-treated (HIV^+^ART^+^) hu-BLT mice. The experimental flow chart is depicted in Figure 1. Briefly, hu-BLT mice were infected with NL4.3-ADA-GFP HIV-1 and, at 6-7 weeks post-infection, a group of mice were treated with ART daily for an additional period of 4-6 weeks. Uninfected hu-BLT mice (HIV^−^) were used as negative controls. Results in Supplemental Figure 1A-B depict the typical engraftment of human cells (lacking the expression of mCD45) in the liver and lungs of HIV^−^, HIV^+^, and HIV^+^ART^+^ BLT mice. Human CD3^+^ T-cells expressing or not CD4 (Supplemental Figure 1C-D), as well as CD3^−^CD33^+^ myeloid cells (subsequently referred to as MΦ) (Supplemental Figure 1E-F), were present in these tissues at frequencies that allowed for their further sorting by flow cytometry for HIV-1 reservoir studies.

**Figure 1:**
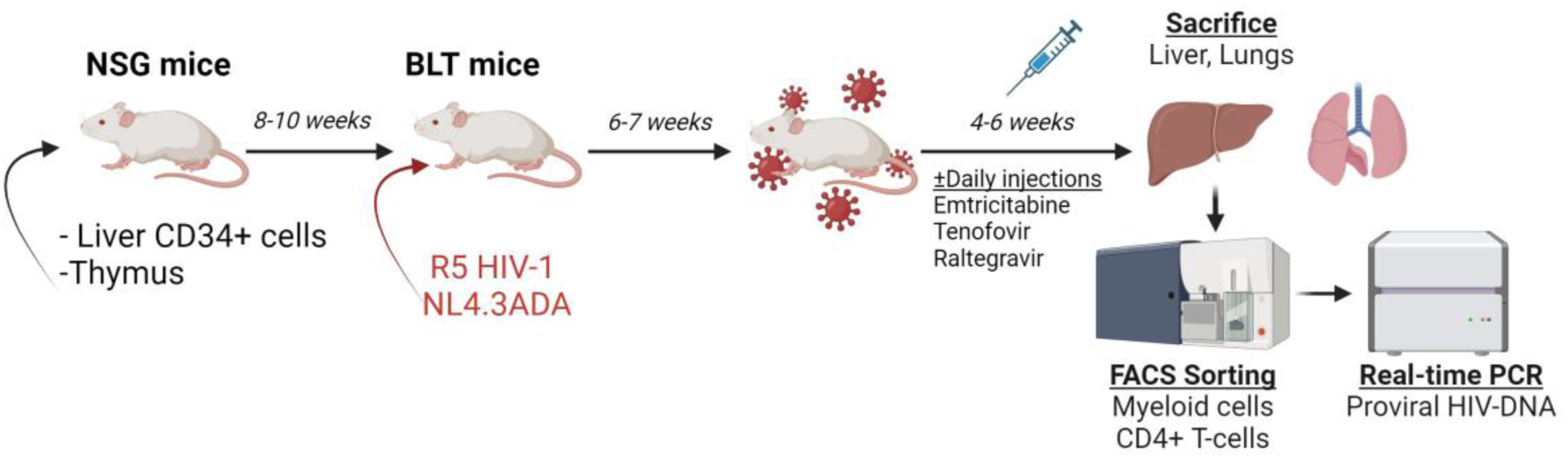
Experimental flowchart for hu-BLT mouse generation, HIV-1 infection and ART. Hu-BLT mice were infected with the CCR5-tropic NL4.3-ADA-GFP HIV-1 *via* intraperitoneal injection. HIV-1 viral load was determined in the plasma of humanized mice weekly using the quantitative COBAS AmpliPrep/COBAS TaqMan HIV-1 test, version 2.0, Roche (detection limit, <20 copies/ml). The ART treatment included a combination of emtricitabine (100 mg/kg weight), tenofovir (50 mg/kg), and raltegravir (68 mg/kg). For a group of HIV-infected hu-BLT mice, ART was administered daily at 6-7 weeks post-infection (wpi) and maintained for 4-6 weeks until sacrifice (HIV^+^ART^+^). A control HIV-infected group remained ART-naive for the duration of the experiment (HIV^+^ART^−^).

### Macrophages residing in the liver support integrative HIV-1 infection

The liver is an important lymphoid organ, containing embryonically derived Kupffer cells, as well as BM monocyte-derived MΦ (16), with the highest CD14 expression documented on monocyte derived SL-MΦ (30–32). In the first set of experiments, liver-infiltrating cells isolated from HIV^+^ART^−^ hu-BLT mice (<u>Batch 1 – Donor A</u>) were stained with mCD45 Abs to identify mouse immune cells, as well as with a cocktail of human Abs to identify human T-cells (CD3 and CD4) and myeloid subsets (HLA-DR, CD68, CD33, CD14). Matched CD14^+^ and CD14^−^ MΦ, co-expressing HLA-DR, CD68 and CD33, and CD4^+^ T-cells, were sorted by flow cytometry for integrated HIV-DNA quantification (Figure 2A). Both CD14^+^ and CD14^−^ MΦ showed permissiveness to integrative infection, with CD14^−^ MΦ (mean: 1,987 copies/10^6^ cells) carrying higher levels of proviral HIV-DNA compared to CD14^+^ MΦ (mean: 348 copies/10^6^ cells) (Figure 2B). As expected, CD4^+^ T-cells expressed greater levels of integrated HIV-DNA (mean: 12,316 copies/10^6^ cells) compared to MΦ (Figure 2B). Of note, all PCR quantifications were performed on a similar number of cells, as reflected by the CD3 counts (2 CD3 copies/cell) (Figure 2C). These experiments were performed with cells pooled per organ from n=4 mice with detectable plasma viral load at the time of sacrifice (range: 3.4×10^5^ to 5.9×10^5^ HIV-RNA copies/mL) (Figure 2D).

**Figure 2:**
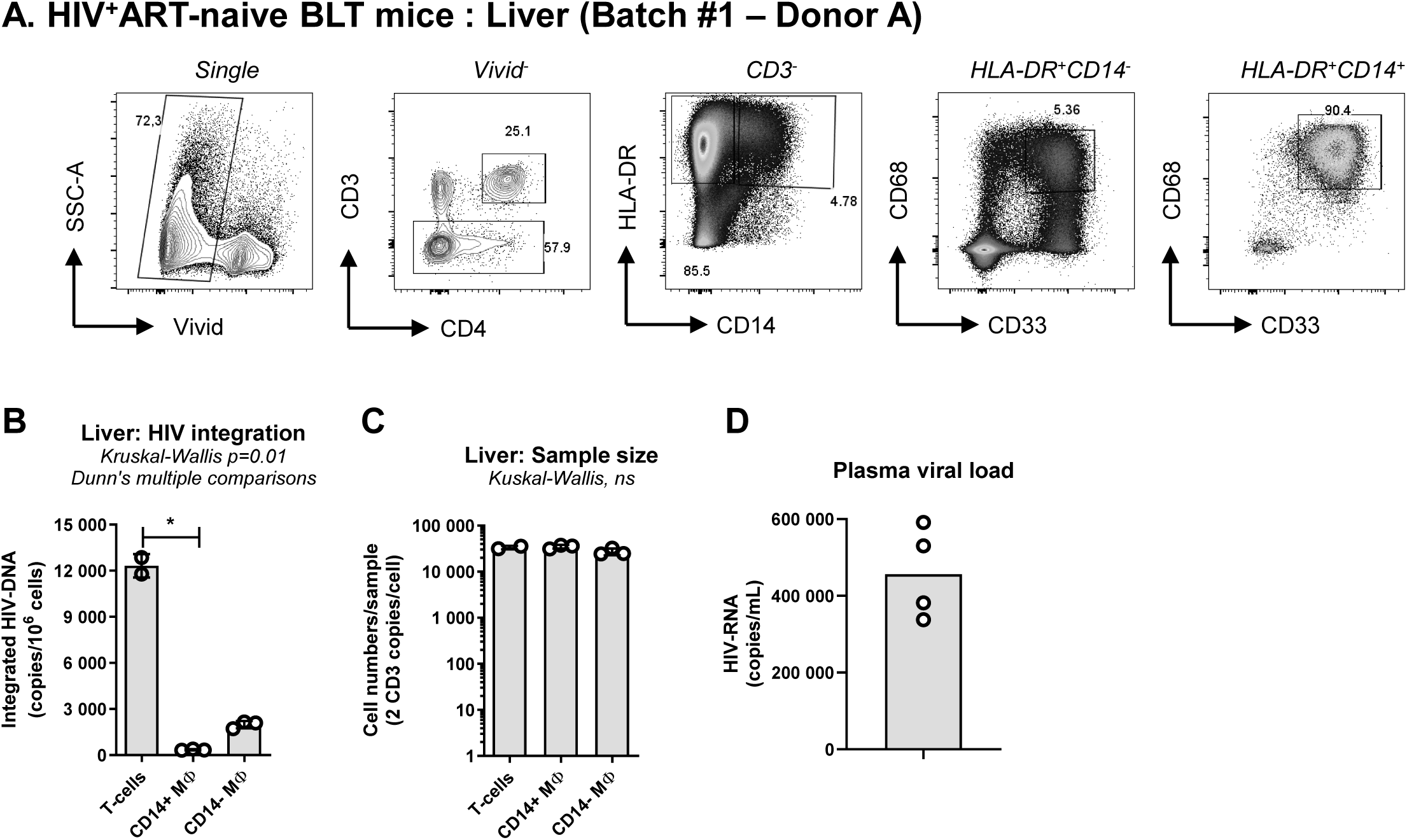
Integrative HIV-1 infection in liver CD14^+^ and CD14^−^ MΦ of HIV-infected ART-naive hu-BLT mice. Cells infiltrating the liver of HIV^+^ hu-BLT mice were isolated, pooled per organ for mice from the same group, and stained with a cocktail of fluorochrome-conjugated Abs. **(A)** Shown is the gating strategy for the identification of CD4^+^ T-cells (identified as CD3^+^CD4^+^) and CD14^+^ *versus* CD14^−^ MΦ (identified as CD3^−^CD4^+/-^HLADR^+^CD33^+^CD68^+^). **(B-C)** T-cells and myeloid cells were simultaneously sorted by flow cytometry for integrated HIV-DNA quantification. Shown are integrated HIV-DNA copies/10^6^ cells (mean ± SD of triplicate wells) **(B)** and the cell numbers/PCR (2 CD3 copies/cell) **(C)**. Shown is the plasma viral load expressed as HIV-RNA copies/mL of mouse plasma (individual mouse and median values) **(D)**. Experiments were performed with pooled cells from n=4 HIV^+^ hu-BLT mice. The hu-BLT mice were made with fetal tissues of Donor A

In the second set of experiments, cells purified from liver tissues of HIV^+^ART^−^ hu-BLT mice (<u>Batch 2 – Donor B</u>) were sorted by FACS using a different gating strategy (Figure 3A). In addition to the myeloid markers CD68 and CD33, the chemokine receptor CCR2, previously used to distinguish between LL-TRM (CCR2^−^) and SL-MΦ (CCR2^+^) (24), was included in the cocktail to potentially distinguish between resident embryonically-derived (Lineage^−^CD3^−^ CD33^+^CD68^+^CCR2^−^) and BM-derived (Lineage^−^CD3^−^CD33^+^CD68^+^CCR2^+^) MΦ. The post-sort purity of CCR2^+^ *versus* CCR2^−^ myeloid cells, identified as mCD45^−^lineage^−^CD3^−^HLA-DR^+^ was >98% (Supplemental Figure 2). Integrated HIV-DNA was again detectable in both MΦ subsets, with slightly higher levels of proviral DNA in CCR2^−^ MΦ (mean: 17,120 copies/10^6^ cells) compared to CCR2^+^ (8,861 copies/10^6^ cells) MΦ, but at lower levels compared to CD4^+^ T-cells (mean: 68,063 copies/10^6^ cells) (Figure 3B). Although similar numbers of CCR2^−^ and CCR2^+^ MΦ were analyzed by PCR for integrated HIV-DNA quantification, the sample size was significantly lower for CCR2^−^ MΦ compared to CD4^+^ T-cells (Figure 3C). All six mice tested exhibited a detectable plasma viral load (range: 3.5×10^5^ to 2.5×10^6^ HIV-RNA copies/mL) (Figure 3D). Thus, MΦ subsets identified based on their differential expression of CD14 or CCR2 support integrative HIV-1 infection in the liver of HIV^+^ART^−^ hu-BLT mice, but at lower levels compared to their CD4^+^ T-cell counterparts.

**Figure 3:**
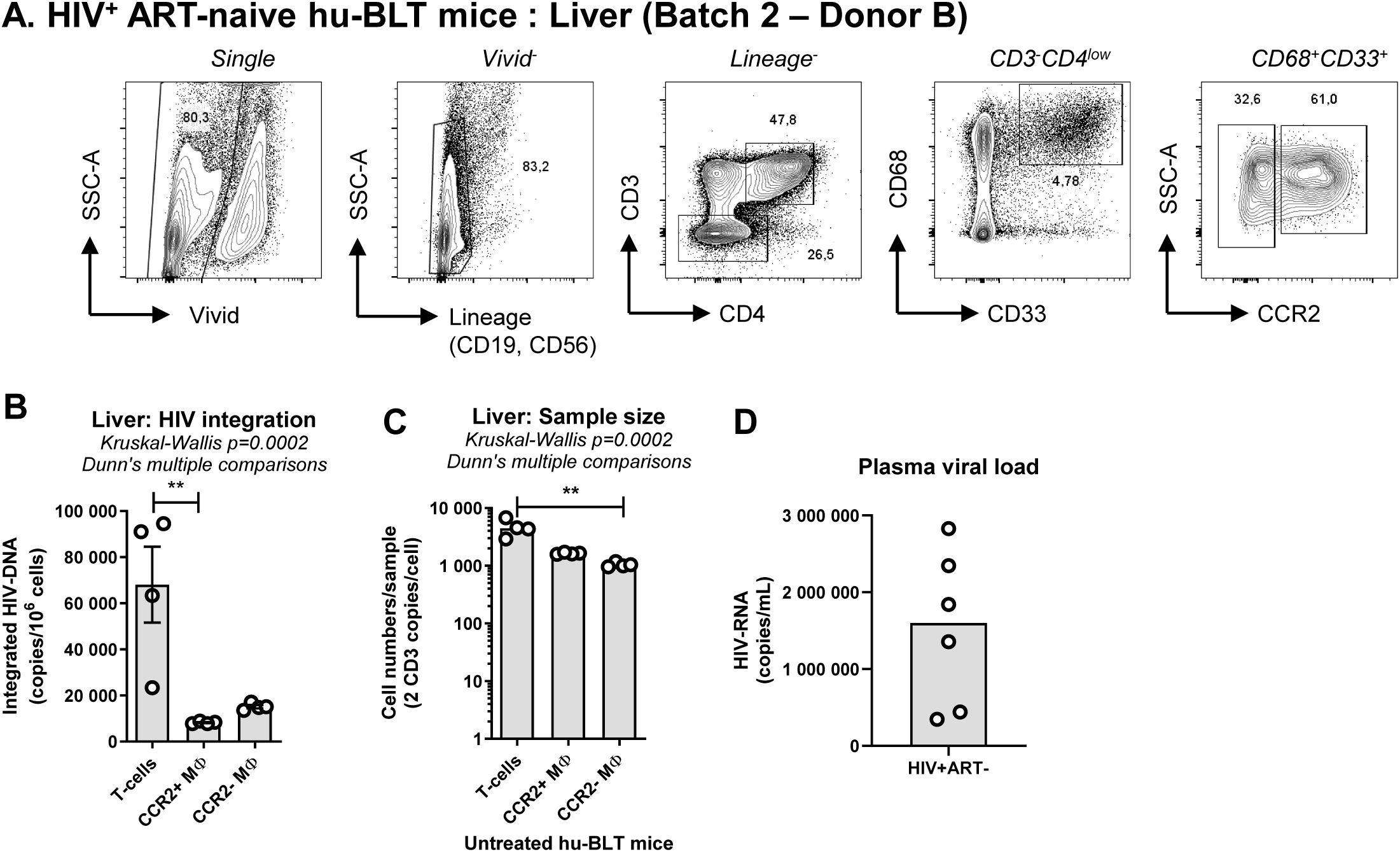
Integrative HIV-1 infection in liver CCR2^+^ and CCR2^−^ MΦ of HIV-infected ART-naive hu-BLT mice. Cells infiltrating the liver of HIV^+^ART^−^ hu-BLT mice were isolated and stained with a cocktail of B cells (CD19) and NK cells (CD56) Abs (Lineage), as well as Abs recognizing T-cells (CD3, CD4) and myeloid cells (CD68, CD33). A viability dye was used to exclude dead cells (Vivid^+^) from the analysis. **(A)** Shown is the gating strategy for the identification of CD4^+^ T-cells (Lineage^−^CD3^+^CD4^+^) and CCR2^+^ *versus* CCR2^−^ MΦ (Lineage^−^CD3^−^ CD4^−/low^CD33^+^CD68^+^). **(B-C)** Liver cells were simultaneously sorted by flow cytometry for integrated HIV-DNA quantification relative to CD3 copies/PCR. Shown are integrated HIV-DNA levels/10^6^ cells **(B)** and cell numbers/PCR (2 CD3 copies/cell) **(C)**. Experiments in **A-C** were performed with pooled cells from n=6 HIV^+^ hu-BLT mice of the same batch (#2). The hu-BLT mice from this Batch were made with fetal tissues from Donor B. Plasma viral load at sacrifice expressed as HIV-RNA copies/mL of plasma are indicated in the graph (individual and median values) **(D)**.

### CD4^+^ T-cells but not MΦ contribute to HIV-DNA persistence during ART in the liver and lungs of hu-BLT mice

In addition to the liver, the lungs are also organs rich in long-lived embryonically derived alveolar MΦ (16, 30). To explore HIV-1 persistence during ART in myeloid cells in this model, cells were isolated in parallel from the liver (Figure 4A-C) and lungs (Figure 4D-F) of HIV^+^ART^+^ hu-BLT mice (<u>Batch 3 – Donor C</u>) and used for flow cytometry sorting of matched CCR2^−^ and CCR2^+^ myeloid cells (mCD45^−^Lineage^−^CD3^−^HLA-DR^+^), as well as CD4^+^ T-cells (mCD45^−^lineage^−^CD3^+^CD4^+^) (Figure 4A and D). In this lot of mice, only a low frequency of cells expressed CD33; therefore, only HLA-DR was used to identify myeloid cells in the liver (Figure 4A) and lungs (Figure 4D). While HIV-DNA integration was detected in CD4^+^ T-cells from both liver (mean: 294 copies/10^6^ cells) and lungs (mean: 298 copies/10^6^ cells), HIV-DNA levels were undetectable in liver CCR2^+^ MΦ and lung CCR2^−^ and CCR2^+^ MΦ of HIV^+^ART^+^ hu-BLT mice (Figure 4B and 4E). It is noteworthy that the number of cells used *per* PCR reaction for CD4^+^ T-cells was markedly higher compared to that of MΦ from liver and lungs, as indicated by the CD3gene quantification (Figure 4C and 4F). Also, the CD3 counts were at similar levels in CCR2^+^ and CCR2^−^ MΦ from the lungs, while in the liver the CD3 gene expression was undetectable in CCR2^−^ MΦ (Figure 4C and 4F), due to the insufficient number of cells sorted by flow cytometry. As expected, mice used in these experiments exhibited undetectable plasma viral load in response to ART (Figure 4G). Thus, these results demonstrate preferential HIV-DNA persistence during ART in CD4^+^ T-cells but not CCR2^+^/CCR2^−^ MΦ isolated from the liver and lungs of hu-BLT mice. However, considering differences in the cell number analyzed between T-cells and MΦ, it is hard to exclude the contribution of MΦ to HIV-1 persistence during ART, especially for liver CCR2^−^ MΦ.

**Figure 4:**
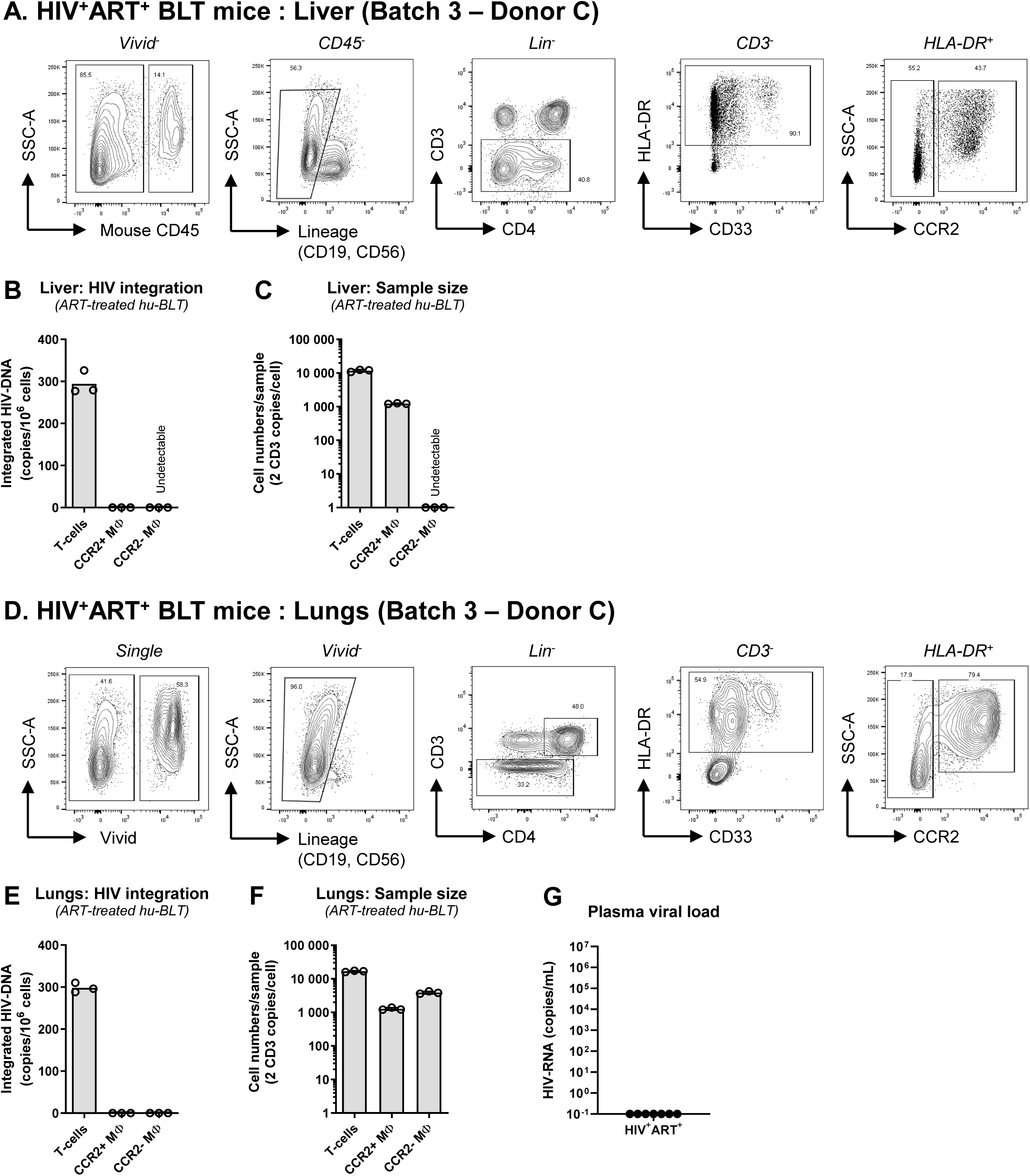
CD4^+^ T-cells but not MΦ carry integrated HIV-DNA in the liver and lungs of HIV-infected ART-treated hu-BLT mice. Cells were isolated from the liver and lungs of HIV^+^ART^+^ hu-BLT mice were stained with a cocktail of B cells (CD19) and NK cells (CD56) Abs (Lineage), as well as Abs recognizing T-cells (CD3, CD4) and myeloid cells (HLA-DR, CD33). A viability dye was used to exclude dead cells (Vivid^+^) from the analysis. The mouse CD45 Ab (mCD45) was used to distinguish between human and mouse immune cells. Shown is the gating strategy for the identification of CD4^+^ T-cells (mCD45^−^Lineage^−^CD3^+^CD4^+^) and CCR2^+^ *versus* CCR2^−^ MΦ (mCD45^−^Lineage^−^CD3^−^HLA-DR^+^CD33^+^) in matched samples from the liver **(A-C)** and lungs **(D-F)**. CD4^+^ T-cells and MΦ were simultaneously sorted by flow cytometry for integrated HIV-DNA quantification in the liver **(B-C)** and lungs **(E-F),** relative to cell counts. Shown are integrated HIV-DNA levels *per* 10^6^ cells **(B and E)** and the cell numbers *per* PCR reaction **(C and F)**. Experiments in **A-C** and **D-F** were performed with pooled cells from n=7 HIV^+^ART^+^ hu-BLT mice of the same organ and batch (#3). These mice were generated with fetal tissues from Donor C. Plasma viral levels expressed as HIV-RNA copies/mL of plasma were quantified in the corresponding mice shown in Panels a-f and depicted in the graph (individual and median values) **(G)**.

To further investigate HIV-1 reservoir persistence during ART, the fourth set of experiments was performed with cells extracted from the liver and lungs of HIV^+^ART^−^ (n=8) and HIV^+^ART^+^ (n=4) hu-BLT mice, and pooled per organ originating from the same lot (<u>Batch 4;</u> Figure 5A-G). The CCR2^−^ and CCR2^+^ myeloid cells (mCD45^−^Lineage^−^CD3^−^HLA-DR^+^CD33^+^CD68^+^), as well as CD4^+^ T-cells (mCD45^−^Lineage^−^CD3^+^CD4^+^) from liver and lungs, were identified as depicted in Figure 5A and 5D, respectively.

**Figure 5:**
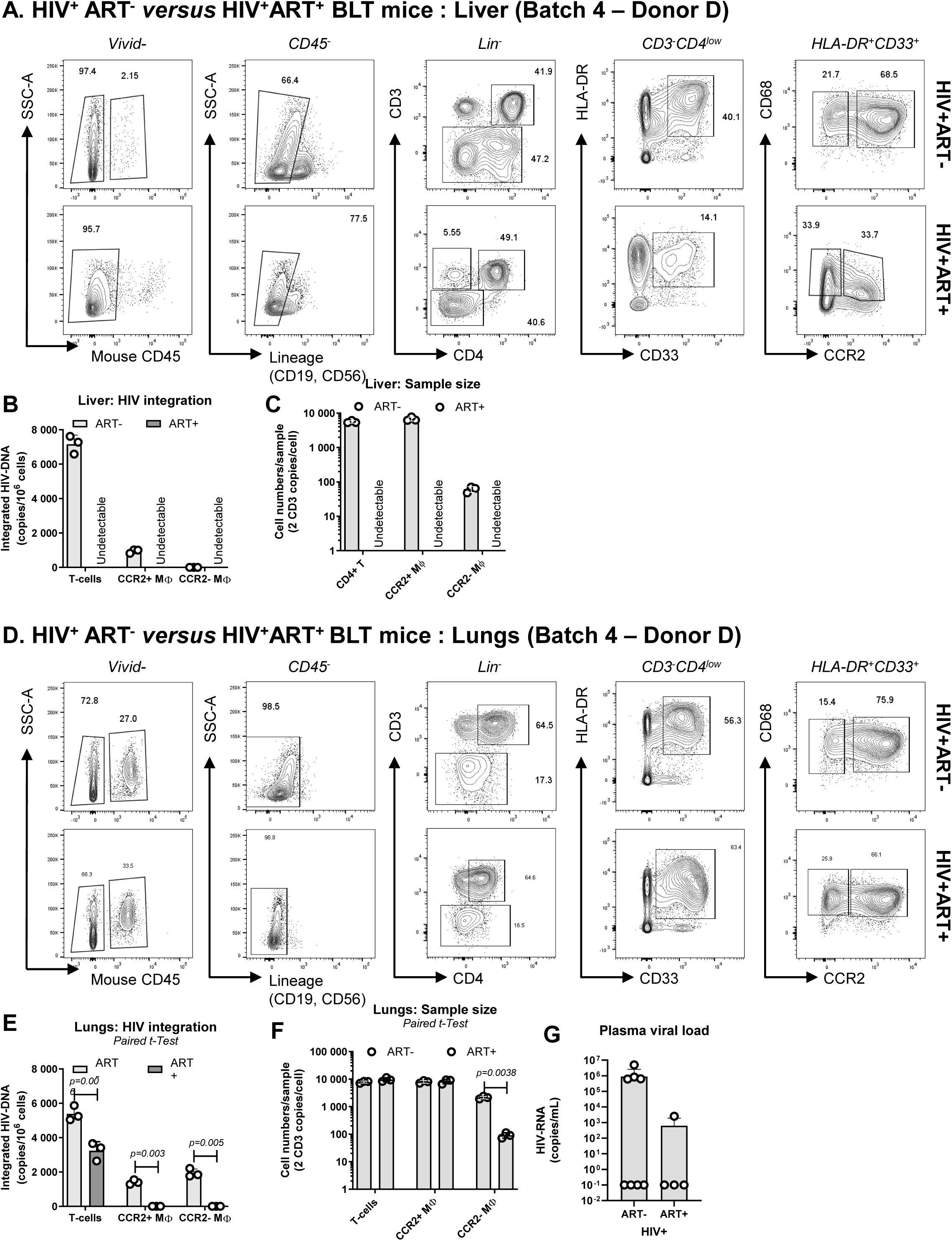
Integrative HIV-1 infection in MΦ is detectable before but not after ART initiation in HIV-infected hu-BLT mice. Cells were isolated from the liver and lungs of HIV^+^ART^−^ and HIV^+^ART^+^ hu-BLT mice from the same organ and batch (#4) and stained with a cocktail of fluorochrome-conjugated Abs **(A-D)**, similar to Figures 1-2. Shown is the gating strategy for the identification of CD4^+^ T-cells (mCD45^−^Lineage^−^CD3^+^CD4^+^) and CCR2^+^ *versus* CCR2^−^ MΦ (mCD45^−^Lineage^−^CD3^−^HLA-DR^+^CD33^+^CD68^+^) in the liver **(A)** and lungs **(D)** of HIV^+^ART^−^ **(top panels)** and HIV^+^ART^+^ **(bottom panels)** hu-BLT mice. Cells were simultaneously sorted by flow cytometry for integrated HIV-DNA quantification in CD4^+^ T-cells and MΦ from the liver **(B-C)** and lungs **(E-F)** by real-time nested PCR, upon normalization to CD3 counts. Shown are levels of integrated HIV-DNA copies/10^6^ cells **(B and E)** and the cell numbers/PCR reactions **(C and F)**. The experiment was performed with pooled cells from n=8 HIV^+^ART^−^ and n=4 HIV^+^ART^+^ hu-BLT mice from the same organ and batch (#4). Hu-BLT mice were made from fetal tissues of Donor D. Plasma viral load levels expressed as HIV-RNA copies/mL are depicted in the graph (individual and median values) **(G)**.

In the liver, the frequency of CD4^+^ T-cells was similar in ART-naive (ART^−^) and ART-treated (ART^+^) HIV^+^ hu-BLT mice (Figure 5A). However, the frequency of HLA-DR^+^CD33^+^ myeloid cells was decreased in HIV^+^ART^+^ *versus* HIV^+^ART^−^ hu-BLT mice (40% *versus* 14% of CD3-cells) (Figure 5A). In HIV^+^ART^−^ hu-BLT mice, all HLA-DR^+^CD33^+^ myeloid cells expressed CD68; while in HIV^+^ART^+^ hu-BLT there was a downregulation of CD68 expression (Figure 5A). In the liver of HIV^+^ART^−^ hu-BLT mice, HIV-DNA integration was detected at the highest levels in CD4^+^ T-cells (7,158 copies/10^6^ cells) and at lower levels in CCR2^+^ MΦ (947 copies/10^6^ cells), while HIV-DNA levels were undetectable in CCR2^−^ MΦ (Figure 5B). The PCR amplifications in HIV^+^ART^−^ hu-BLT mice were performed for CD4^+^ T-cells and CCR2^+^ MΦ on similar cell numbers/reactions; however, the number of cells sorted was reduced for CCR2^−^ MΦ (Figure 5C), thus preventing us from drawing a definite conclusion. Finally, in this lot, the number of cells extracted from the liver of HIV^+^ART^+^ hu-BLT mice was too low to allow for the sorting of sufficient numbers of myeloid cells for accurate quantification of integrated HIV-DNA (Figure 5B-C).

In the lungs, the phenotype of myeloid cells from HIV^+^ART^−^ and HIV^+^ART^+^ hu-BLT mice was similar (Figure 5D). HIV-DNA integration was detected in CD4^+^ T-cells of HIV^+^ART^+^ (3,235 copies/10^6^ cells) but at lower levels compared to ART^−^ (5,386 copies/10^6^ cells) hu-BLT mice (Figure 5E). Also, integrated HIV-DNA was detected in both CCR2^+^ (1,406 copies/10^6^ cells) and CCR2^−^ (1,939 copies/10^6^ cells) MΦ of HIV^+^ART^−^, but not in CCR2^+^/CCR2^−^ MΦ of HIV^+^ART^+^ hu-BLT mice (Figure 5E). The cell number *per* PCR reaction was similarly high for CD4^+^ T-cells and CCR2^+^ MΦ isolated from the lungs of both HIV^+^ART^−^ and HIV^+^ART^+^ hu-BLT mice, but was significantly lower for CCR2^−^ MΦ of HIV^+^ART^+^ *versus* HIV^+^ART^−^ hu-BLT mice (Figure 5F). In this lot of mice, only 4/8 of ART^−^ hu-BLT mice were viremic (range: 6.2×10^5^ to 5.0×10^6^ HIV-RNA copies/mL), whereas 1/4 HIV^+^ART^+^ hu-BLT mice had detectable plasma viral load despite ART (2,520 HIV-RNA copies/mL) (Figure 5G). Experiments were performed with cells pooled from each group of hu-BLT mice.

Together, these results demonstrate that CCR2^+^ and CCR2^−^ MΦ from the liver and lungs are permissive to integrative infection, but do not carry detectable HIV-1 reservoirs in HIV^+^ART^+^ hu-BLT mice. This is in contrast to CD4^+^ T-cells, which are highly permissive to HIV-1 infection in the absence of ART and carry relatively high levels of proviral HIV-DNA during ART.

### Human MΦ with a LL-TRM phenotype are observed in the liver of hu-BLT mice

LL-TRM in the human liver are defined as CD68^+^CD14^+^ cells expressing markers such as Tim4, CD163, and CX3CR1 (27, 33, 61), while LL-TRM in the myocardium exhibit a CCR2^−^ phenotype and are exclusively replenished through local proliferation of myeloid precursors (24). The development of LL-TRMs in hu-BLT mouse models remains undefined. To address this question, we performed an extensive phenotypic characterization of myeloid cells in the liver and lungs of HIV^−^ hu-BLT mice (n=7, Batch 5 – Donor E). The markers used in Figure 6-7 were reported in the literature to be preferentially expressed on embryonically derived LL-TRM *versus* monocyte-derived SL-MΦ (Supplemental Table 1).

**Figure 6:**
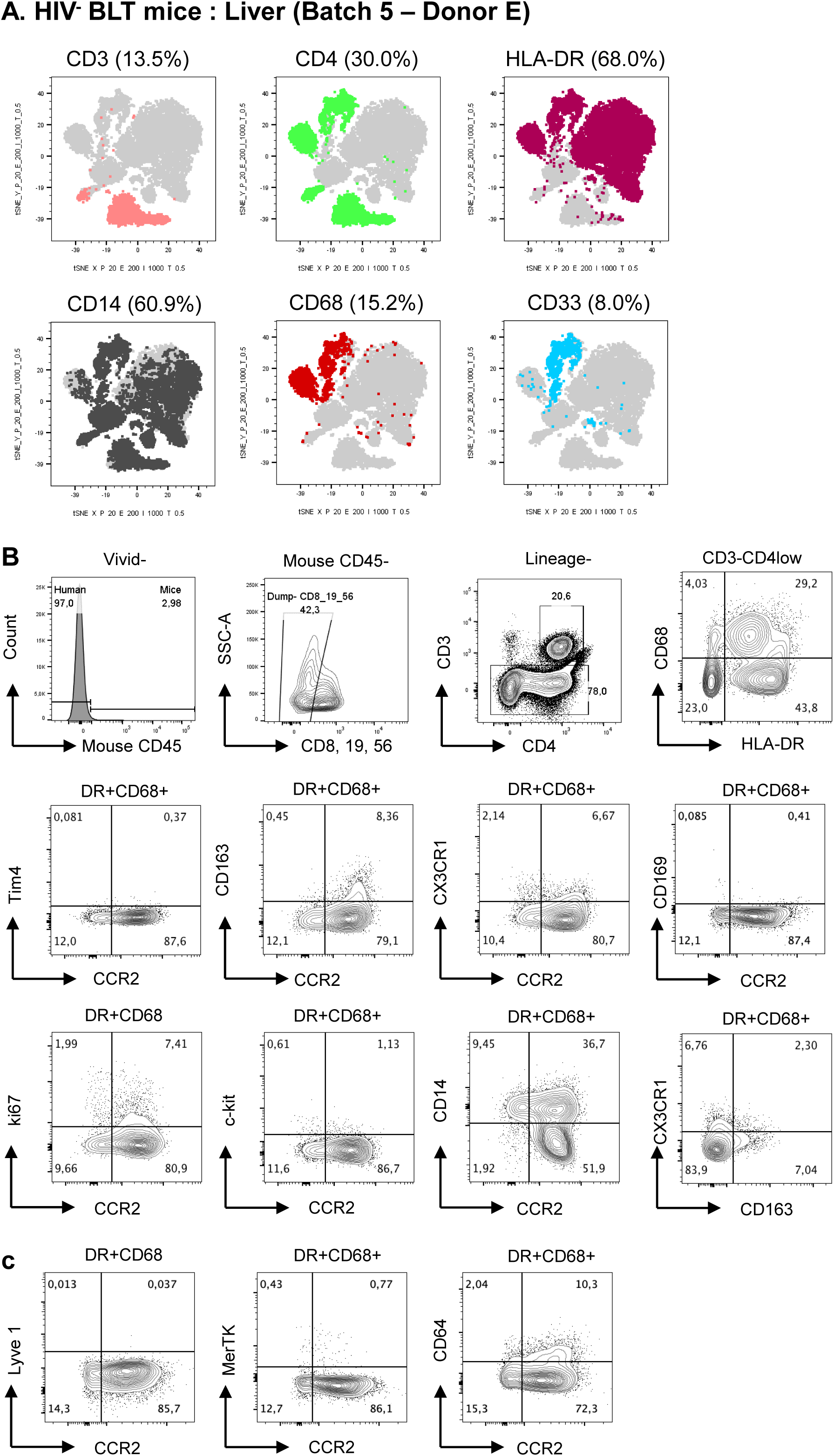
Expression of LL-TRM markers on myeloid cells from the liver of uninfected hu-BLT mice. Cells from the liver of n=7 uninfected hu-BLT mice were mechanically and enzymatically isolated and infiltrating immune cells were stained with a cocktail of human Abs recognizing CD3, CD4, CD8, CD56, CD19, CD45RA, CD14, CD68, HLA-DR, CD33, CCR2, CX3CR1, c-Kit, CD163, Ki67, Tim4, CD64, Lyve1, and MerTK, as well as mouse CD45 Abs. Dead cells were excluded using the LIVE/DEAD cell stain kit. Cells were analyzed by flow cytometry (BD LSRII). tSNE plots were generated using Flow Jo V10.5.2. Shown is the distribution of myeloid markers in the lineage-negative population (CD19^−^CD56^−^CD8^−^) in the liver **(A)**. Shown is the expression of known LL-TRM markers on CD68^+^HLA-DR^+^ MΦ and their co-expression with CCR2 in the liver **(B-C)**. Experiments are performed with pooled cells from uninfected mice, which were made from fetal tissues of Donor E.

**Figure 7:**
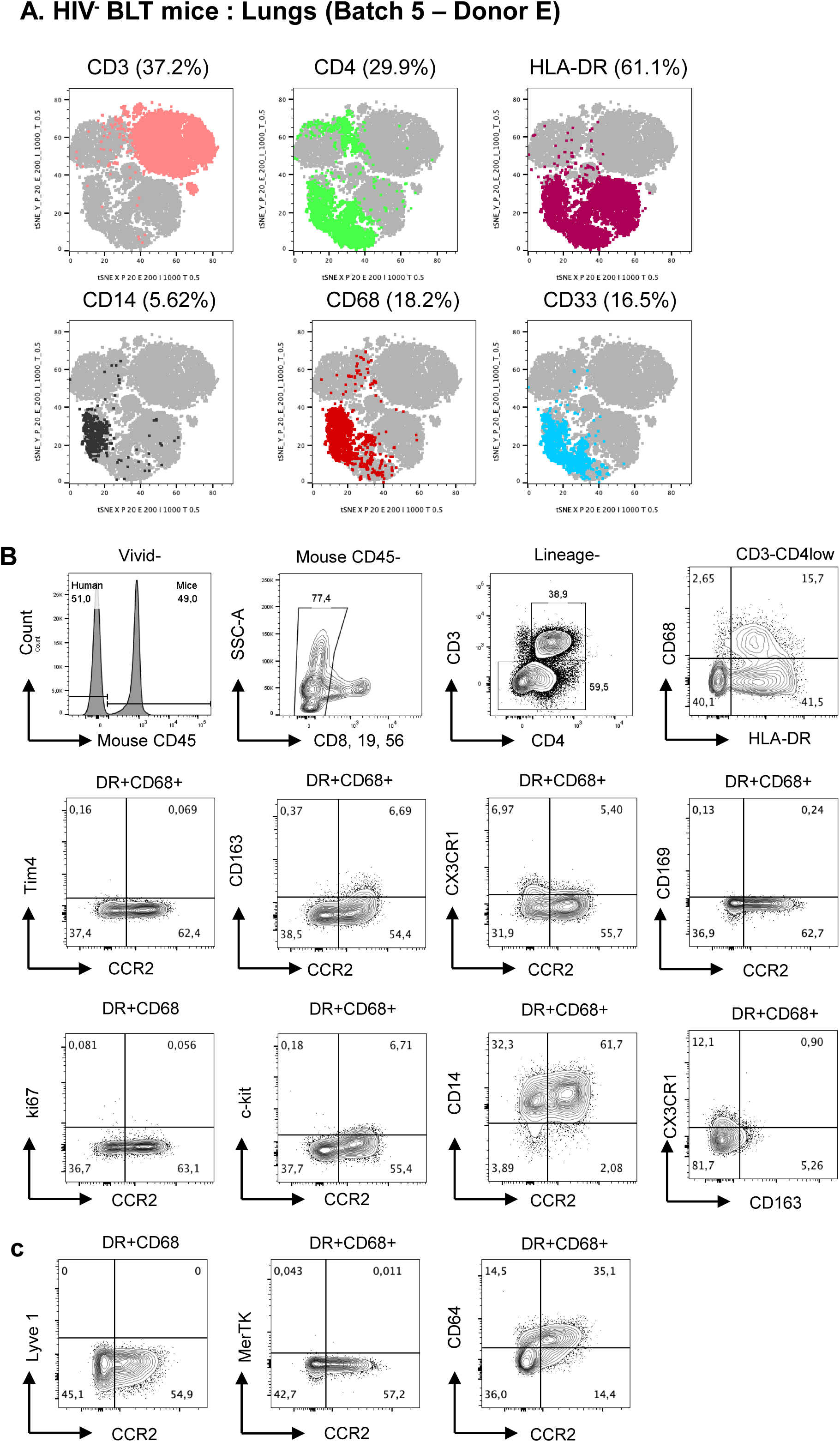
Expression of LL-TRM markers on myeloid cells from the lungs of uninfected hu-BLT Mice. Cells from the lungs of n=7 uninfected hu-BLT mice were mechanically and enzymatically extracted and infiltrating immune cells were stained with a cocktail of human Abs as in Figure 6. Dead cells were excluded using the LIVE/DEAD cell stain kit. Cells were analyzed by flow cytometry (BD LSRII). tSNE plots were generated using Flow Jo V10.5.2. Shown is the distribution of myeloid markers in the lineage-negative population (CD19^−^CD56^−^CD8^−^) in the lungs **(A)**. Shown is the expression of known LL-TRM markers in CD68^+^HLA-DR^+^ MΦ and their co-expression with CCR2 in the lungs **(B-C)**. Experiments are performed with pooled cells from uninfected mice, which were made from fetal tissues of Donor E.

In the <u>liver</u> of uninfected hu-BLT mice, the differential expression of CD3, CD4, HLA-DR, CD14, CD68, and/or CD33 allowed us to visualize myeloid subsets in tSNE plots (Figure 6A). CD68 expression was observed on a fraction of CD3^−^HLA-DR^+^ cells (<35%), while CD14 was expressed on a fraction of CD3^−^HLA-DR^+^CD68^+^ cells (<50%) (Figure 6B). The LL-TRM markers CD163 and CX3CR1 were principally expressed on a small fraction of CCR2^+^ myeloid cells (CD68^+^HLA-DR^+^), with only few cells co-expressing CD163 and CX3CR1 (<3%) and low/undetectable Tim4 and c-Kit expression (Figure 6B). Interestingly, the surrogate marker of proliferation Ki67 was expressed on a small fraction of CCR2^+^ (7.4%) and to a lesser extent on CCR2^−^ (2%) HLA-DR^+^CD68^+^ myeloid cells (Figure 6B). Other MΦ markers such as MerTK and Lyve1 were not detected on HLA-DR^+^CD68^+^ myeloid cells from the liver, while CD64 was expressed on a small fraction of CCR2^+^ liver MΦ (Figure 6C).

In the <u>lungs</u> of uninfected hu-BLT mice, the tSNE plot representation allowed the visualization of human cells which differentially express CD3, CD4, HLA-DR, CD14, CD68 and/or CD33 (Figure 7A). As in the liver, CD68 expression was observed on a fraction of lung HLA-DR^+^CD3^−^ cells (Figure 7B). In contrast, the majority of CD3^−^HLA-DR^+^CD68^+^ MΦ expressed CD14. Further, CD163 was principally expressed on CCR2^+^ MΦ (<7%), while CX3CR1 was expressed at low levels on both CCR2^+^ and CCR2^−^ cells, with only a few MΦ (<1%) co-expressing the LL-TRM markers CD163 and CX3CR1 (Figure 7B). Other MΦ markers such as Ki67, MerTK and Lyve1 were not expressed on HLA-DR^+^CD68^+^ myeloid cells, while CD64 was expressed on fractions of CCR2+ and CCR2-lung MΦ (Figure 7C).

Together, these results demonstrate the phenotypic heterogeneity of HLA-DR^+^CD68^+^ myeloid cells present in the liver and lungs of the uninfected hu-BLT mice in terms of CD14 expression and reveal the expression of Ki67, a surrogate marker of proliferation, on a fraction of MΦ from the liver, pointing to the fact that a small fraction of LL-TRM can develop in this anatomic site of hu-BLT mice, likely from liver CD34^+^ precursors.

## DISCUSSION

In this study, we employed a hu-BLT mouse model, previously used by our group (56, 57), to explore the contribution of liver and lung MΦ, relative to CD4^+^ T-cells, to HIV-1 reservoir persistence during ART, and to perform an extensive characterization of LL-TRM and SL-MΦ phenotypes of MΦ from these tissues. Our results revealed that MΦ from the liver and lungs, expressing or not the human TRM marker CCR2 (24), were permissive to integrative HIV-1 infection in ART-naive HIV-infected hu-BLT mice. However, levels of proviral HIV-DNA were low/undetectable in these myeloid subsets isolated from the liver and lungs of ART-treated HIV-infected hu-BLT mice. These findings were consistent with the inadequate development of LL-TRM we observed in this hu-BLT model, with the presence of only a small fraction of proliferating (CD163^+^CX3CR1^+^Ki67^+^) MΦ being detected in the liver but not the lungs. This small fraction of LL-TRM likely developed from CD34^+^ precursors uniquely in the liver environment. Thus, although tissue-resident MΦ develop properly in the lungs and liver of hu-BLT mice and are highly permissive to HIV-1 infection, their contribution to HIV-1 reservoir persistence during ART is negligible, likely due to their short-survival and increased turnover *in vivo*.

Models of humanized mice were historically developed for HIV-1 studies and are now used for multiple research purposes including drug development in virology, cancer, or transplantation research (52, 54, 55, 62–64). Different models of humanized mice are currently available, with each model having a specific phenotype and characteristics (65). At the time of the study design, the literature documented CD14 as a marker of Kupffer cells in the liver (27, 61), and CCR2 as a marker distinguishing between LL-TRM (CCR2^−^) and SL-MΦ (CCR2^+^) in the human heart (24). Using a combination of well-established myeloid markers (*i.e.,* HLA-DR, CD68, CD33), we sorted by flow cytometry highly pure CD14^+^ *versus* CD14^−^ and CCR2^+^ *versus* CCR2^−^ MΦ from the liver and lungs. Matched CD4^+^ T-cells from the same organs were sorted in parallel as controls. In the absence of ART, we demonstrated the presence of integrated HIV-DNA in MΦ from HIV-infected hu-BLT mice, and this at slightly higher levels in CD14^−^ and CCR2^−^ MΦ compared to their CD14^+^ and CCR2^+^ counterparts, respectively. Compared to myeloid cells, CD4^+^ T-cells carried higher levels of integrated HIV-DNA in ART-naive hu-BLT mice. In ART-treated hu-BLT mice, integrative infection was low/undetectable in all myeloid subsets sorted. Nevertheless, integrated HIV-DNA levels remained relatively high in CD4^+^ T-cells isolated form the lungs and liver of ART-treated hu-BLT mice, thus validating the persistence of viral reservoir during ART in this model.

Our inability to detect integrative infection during ART in myeloid cells, even when sufficient numbers of cells were sorted, may be explained by the rapid turnover of these cells *in vivo*, pointing to the inappropriate development of long-lived myeloid cells in this model. To explore this possibility, we performed an extensive phenotypic characterization of myeloid cells. Based on our results, this hu-BLT mouse model appears to mainly develop SL-MΦ, which are typically derived from monocytes, to the detriment of LL-TRM of embryonic origin. Markers such as Tim4, CD169, CD163, or CX3CR1 were reported as preferentially expressed by LL-TRM in mice (66, 67). In our model, only a small fraction of MΦ in the liver but not the lungs expressed the LL-TRM markers CD163 and CX3CR1, and the surrogate proliferation marker Ki67. The development of LL-TRM in humanized mice theoretically depends on the existence of appropriate embryonic precursors from human abortive tissues, the existence of proper tissue-specific niches, and the presence of human growth factors (68). In the hu-BLT model used in this study, the NSG mice were engrafted with fetal thymic tissue under the kidney capsule and injected with autologous hepatic CD34^+^ hematopoietic stem cells (HSCs), as previously described (57). The liver CD34^+^ HSCs typically give rise to the third wave of hematopoiesis, leading to the appearance of BM-derived MΦ(16). Among other existent humanized mouse models with improved monocytes/MΦ development, the IL-3/GM-CSF knock-in model in a Rag2^−/-^Il2rγ^−/-^ background allows for the development of alveolar MΦ in the lungs (69), while the MISTRG (M-CSF^h/h^IL-3/GM-CSF^h/h^SIRPa^h/h^TPO^h/h^RAG2^−/-^IL2Rg^−/-^) (70) model enables the development of tissue MΦ in the liver and lungs (71, 72). In addition, the lung-only and hu-BLT spleen humanized mouse models were created to improve and overcome the limitation of other models in terms of tissue-specific human cell differentiation (73). However, whether tissues MΦ with *bona fide* LL-TRM features could indeed be developed in these models remains unclear.

In the liver of uninfected hu-BLT mice, only 15% of the myeloid cells were CD68^+^, which is consistent with other published studies on healthy human liver (61). A recent study using mismatched liver transplants, comparing donor *versus* recipient cells up to 11 years post-transplant, identified a specific phenotype for LL-TRM and SL-MΦ (33). It revealed that CCR2 expression could not discriminate between the two subsets and that liver resident Kupffer cells expressed CD68 and CD14 and the majority co-expressed CX3CR1 and CD163 (33). These Kupffer cells also expressed CD206 (33). Using a similar gating strategy, we found that only around 8% of the liver HLA-DR^+^CD68^+^ cells co-expressed CD163, CX3CR1 and/or Ki67, suggesting that the hu-BLT mouse model develops only a small fraction of hepatic LL-TRM. The expression of Tim4, a reported marker of LL-TRM from different tissues (74, 75), including the liver (76), was low/undetectable in our current hu-BLT mouse model. The low expression of Tim4 on myeloid cells in our model suggests that these cells may not be Kupffer cells. Also, the co-expression of CD163, CX3CR1 and/or Ki67 with CCR2 suggests that these cells have more of a migratory profile, in contrast to the knowledge that Kupffer cells are resident non-migratory cells and have low CCR2 expression (77).

In addition to the embryonic origin of Kupffer cells, BM-derived monocyte have been shown to give rise to self-renewing and fully differentiated Kupffer-like cells in the liver. This mainly happens in condition where embryonically-derived Kupffer cells are depleted, thus rendering the liver niche available for new BM-derived monocyte infiltration (76). Indeed, the niche and environmental signals play an important role in MΦ development. Therefore, even though our polychromatic flow cytometry analysis points to the existence of a small fraction of LL-TRM, with a CD163^+^CX3CR1^+^Ki67^+^ phenotype in the liver of hu-BLT mice, these cells may most likely originate from monocytes, in an attempt to establish a new niche of Kupffer-like resident MΦ in the humanized mouse liver. Their low expression of Tim4 is consistent with the latter scenario, as Kupffer cells are reported to exhibit a Tim4^+^ phenotype and represent the majority of MΦ in the liver under homeostatic conditions(78). To better characterize tissue-resident MΦ from the liver in hu-BLT mice, additional markers such as MARCO should be considered in future studies. Indeed, MARCO is expressed in Kupffer cells which are populated in the periportal areas of the liver (61).

In the lungs, resident alveolar MΦ have been described as cells exhibiting a CD45^+^HLA-DR^+^CD206^+^CD169^+^CD14^−^CD64^+^ phenotype and are enriched in bronchoalveolar lavages (BAL) compared to digested lung tissues (32). In our study, lung tissues were enzymatically and mechanically digested (57). This may explain in part the low frequency of alveolar MΦ with a LL-TRM phenotype in hu-BLT mice. A comparison of BAL *versus* digested tissue will be required to validate the presence of resident alveolar MΦ cells in this system.

## LIMITATIONS OF THE STUDY

Our study has several limitations. First, we were unable to sort sufficient numbers of CCR2^+^/CCR2^−^ MΦ from the liver of HIV^+^ART^+^ hu-BLT mice for viral reservoir studies. Thus, we cannot exclude the possibility that these cells carry viral reservoirs during ART, especially since we observed that liver MΦ express Ki67 and may represent a cells with self-renewal capacity. Future studies are needed to assess HIV-1 reservoir persistence in Ki67^+^ MΦ from the liver. However, we succeeded in sorting CCR2^+^/CCR2^−^ MΦ from both liver and lungs of HIV^+^ART^−^ hu-BLT mice and concluded on their permissiveness to integrative HIV-1 infection *in vivo*. In addition, we successfully sorted CCR2^+^ and CCR2^−^ MΦ from the lungs of HIV^+^ART^+^ hu-BLT mice and observed undetectable HIV-1 integration, in contrast to CD4+ T-cells that were harboring viral reservoirs during ART in both the liver and lungs of hu-BLT mice. Nevertheless, we cannot exclude the possibility that HIV-1 reservoirs persist during ART in MΦ from tissues other that the lungs, *e.g.,* spleen and bone marrow, tissues that were not available for this study. Second, since this study used only polychromatic flow cytometry to phenotypically characterize myeloid cells, one cannot exclude the possibility that some surface markers are stripped following enzymatic treatment of tissues. Histology studies represent an important alternative approach for the *in situ* characterization of tissue MΦ. Moreover, single-cell RNA sequencing should be performed to understand whether the protocol used to generate hu-BLT mice has an impact on the development of LL-TRM in various tissues. Furthermore, single-cell RNA sequencing data generated in hu-BLT mouse models should be further compared to public transcriptional signatures available in human TRM. The LL-TRM reconstitution needs to be validated organ by organ given the known tissue-specific MΦ heterogeneity. This will require a combined approach using flow cytometry, histology, and single-cell RNA, as well as a comparison to public human datasets. This will allow the validation of both gene signatures and spatial localization of various myeloid cells in human tissues. Finally, the use of the HIV-RNA/DNA Scope technology (79) on hu-BLT mice and human autopsy tissues, in combination with specific myeloid cell and T-cell markers would further our understanding of HIV reservoir persistence in CD4^+^ T-cells *versus* myeloid cells.

## CONCLUSIONS

In conclusion, while current hu-BLT mouse models represent an important tool for HIV research allowing to perform preclinical testing for new HIV-1 cure interventions, they might not be the best models to study the contribution of myeloid cells to HIV-1 reservoir persistence during ART, unless an optimal development of LL-TRM is achieved. Also, while LL-TRM *versus* SL-MΦ are well characterized in mice (16–18), new studies are needed to identify appropriate surface and functional markers specifically expressed by these MΦ subsets in humans. Such knowledge will be essential to document viral reservoir persistence during ART in human samples, including autopsy tissues (80), as in most recent HIV studies (2, 43, 81, 82). Finally, given the permissiveness to integrative HIV-1 infection of lung and liver MΦ in the absence of ART, and the likelihood that these cells contribute to viral rebound from CD4^+^ T-cells upon ART interruption, HIV-1 cure interventions should target HIV-1 replication in both CD4^+^ T-cells and MΦ to achieve the ultimate goal of viral eradication.

## DECLARATIONS

### Ethics approval and consent to participate

All experiments performed were approved and conformed to the regulatory standards set by the Institutional Review Board (IRB) of the Research Ethics Committees (*CER, Comité d’éthique de la recherche*). Briefly, human fetal tissues were obtained following written informed consent from donors and approval of a research protocol by the Centre Hospitalier Universitaire (CHU) Sainte-Justine IRB (CER# 2126) and CSSS Jeanne-Mance review boards (Montreal, Canada). As per the ethics guidelines, no information regarding the donors was provided. Mice were maintained in a specific pathogen-free animal facility of CHU Sainte-Justine Research Center and in the animal core facility of the Montreal Clinical Research Institute (IRCM) following protocols approved by each institution’s Institutional Animal Care and Use Committee (IRCM 2018-11; IRCM 2015-11&2018-06 and CIBPAR#2021-2961). Experiments on HIV-infected human cells were approved by the Centre de recherche du CHUM institutional review board (CER 2015-5494; CE 14.077-CA) and performed in accordance with the principles of the Declaration of Helsinki. All experiments were performed in accordance with the ARRIVE guidelines and regulations (https://arriveguidelines.org/arrive-guidelines)(83).

### Consent for publication

All study participants consented with the publication of the results

### Availability of data and materials

Data in this manuscript is available upon request to the corresponding author

### Authors’ contributions

AC, TNQP and JD designed and performed research, analyzed data, prepared figures, and wrote the manuscript. NFR and LRM performed research related to mouse tissue processing and flow cytometry cell sorting. OV and TNQP performed HIV infections, ART administration in mice, and mouse tissue processing. J.V.G., N.P., YL, and KB contributed to the production of hu-BLT mice. MAJ and JE contributed to study design and provided protocols for the isolation of immune cells from tissues and the identification of myeloid subsets. E.H. supervised the generation of humanized mice and provided expertise and input regarding the conception of the study. EAC conceived study protocols, contributed to research design, and manuscript writing, and obtained funding. PA conceived the research study hypothesis, designed research, analyzed data, wrote the manuscript, and obtained funding. All co-authors revised the manuscript.

## Acknowledgements

The authors thank Dr. Dominique Gauchat and Philippe St Onge (Flow Cytometry Core Facility, CHUM-Research Center, Montréal, QC, Canada) for expert technical support with polychromatic flow cytometry sorting; Olfa Debbeche (Biosafety Level 3 Core Facility CHUM-Research Center, Montréal, QC, Canada); and Maître Eliane Brunello (Legal Affairs Office, CHUM-Research Center, Montréal, QC, Canada) Mario Legault (FRQ-S/AIDS and Infectious Diseases Network; Montréal, QC, Canada). The authors also appreciate the technical support provided by Fréderic Dallaire as well as by the staff of the animal facilities of the IRCM and CRA-CHU Ste Justine and the IRCM cytometry platform. We also thank Renée Dicaire for her assistance in the generation of hu-BLT mice. The authors would like to thank Dr. Romas Geleziunas at Gilead Sciences for the gift of Emtricitabine and Tenofovir and Dr. Daria Hazuda at Merck for providing us with the Raltegravir.

## SUPPLEMENTAL TABLES

**Supplemental Table 1:** Antibodies used for flow cytometry analysis and sorting.

| **Abs** | **Fluorochromes** | **Clones** | **Source** |
| --- | --- | --- | --- |
| Mouse CD45 | PerCP-Cy5.5 | 30 F11 | Biolegend |
| Hu CD3 | Pacific Blue | UCHT1 | BD |
| Hu CD4 | BUV496 | SK3 | BD |
| Hu CD45RA | BUV737 | HI100 | BD |
| Hu CD8 | FITC | BW135/80 | Miltenyi |
| Hu CD19 | FITC | LT19 | Miltenyi |
| Hu CD56 | FITC | HCD56 | Biolegend |
| Hu CD14 | Alexa Fluor700 | M5E2 | BD |
| Hu CD68 | PE-Cy7 | Y1/82A | Biolegend |
| Hu HLA-DR | BV786 | L243 | Biolegend |
| Hu CD33 | PE | WM53 | Biolegend |
| Hu CCR2 | PE-CF594 | K036C2 | Biolegend |
| Hu CX3CR1 | BV650 | 2A9-1 | Biolegend |
| Hu c-kit | BV605 | 104 D2 | Biolegend |
| Hu CD163 | BV711 | GHI/61 | Biolegend |
| Hu ki67 | BUV395 | B56 | BD |
| Hu Tim4 | APC | 9F4 | Biolegend |
| Hu CD169 | APC-vio770 | 7-239 | Miltenyi |
| Hu CD64 | BV605 | 10.1 | Biolegend |
| Hu CD115 | BV711 | 9-4D2-1E4 | Biolegend |
| Hu CD44 | BUV395 | 515 | BD |
| Lyve-1 | APC | 537028 | R&D |
| MerTK | APC Fire 750 | 590H11G1E3 | Biolegend |

