## Supplementary figures and images for "Humanized Bone Marrow-Liver-Thymus Mice for Studying HIV-1 Persistence in Liver and Lung CD4+ T and Myeloid Cell Subsets during Antiretroviral Therapy"

### Supplemental Figure 1-2

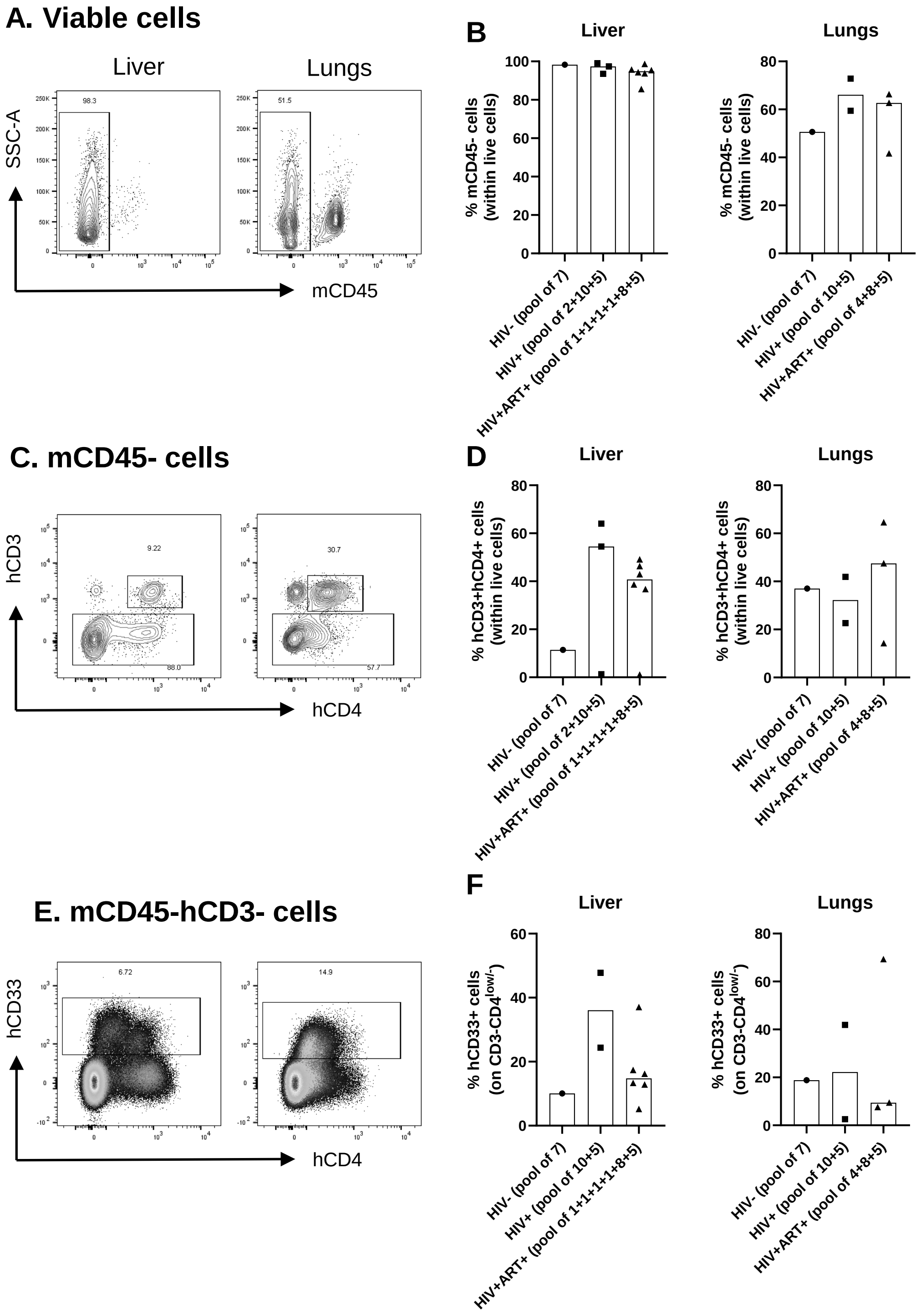


Cattin *et al.*, Supplemental Figure 1

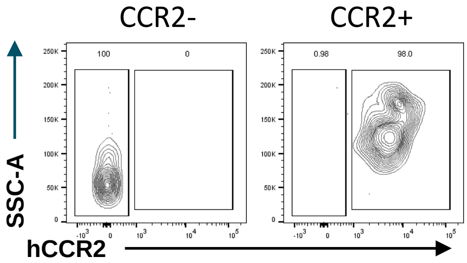
